# Structural and biochemical characterization establishes a detailed understanding of KEAP1-CUL3 complex assembly

**DOI:** 10.1101/2023.02.15.528651

**Authors:** Roslin J Adamson, N Connor Payne, Sergio G. Bartual, Ralph Mazitschek, Alex N Bullock

## Abstract

KEAP1 promotes the ubiquitin-dependent degradation of NRF2 by assembling into a CUL3-dependent ubiquitin ligase complex. Oxidative and electrophilic stress inhibit KEAP1 allowing NRF2 to accumulate for transactivation of stress response genes. To date there are no structures of the KEAP1-CUL3 interaction nor binding data to show the contributions of different domains to their binding affinity. We determined a crystal structure of the BTB and 3-box domains of human KEAP1 in complex with the CUL3 N-terminal domain that showed a heterotetrameric assembly with 2:2 stoichiometry. To support the structural data, we developed a versatile TR-FRET-based assay system to profile the binding of BTB-domain-containing proteins to CUL3 and determine the contribution of distinct protein features, revealing the importance of the CUL3 N-terminal extension for high affinity binding. We further provide direct evidence that the investigational drug CDDO does not disrupt the KEAP1-CUL3 interaction, even at high concentrations, but reduces the affinity of KEAP1-CUL3 binding. The TR-FRET-based assay system offers a generalizable platform for profiling this protein class and may form a suitable screening platform for ligands that disrupt these interactions by targeting the BTB or 3-box domains to block E3 ligase function.

**Graphical abstract:** 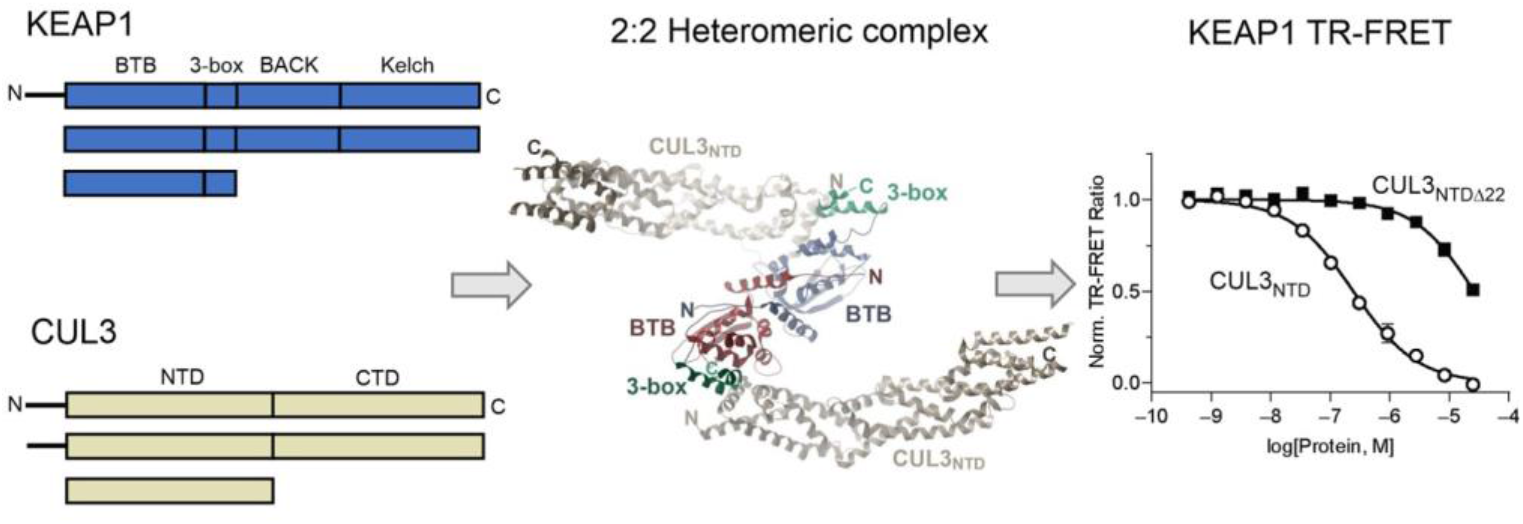

**Highlights:**

- A new crystal structure defines KEAP1 BTB and 3-box domain interactions with CUL3
- KEAP1 and CUL3 form a heteromeric 2:2 complex with a *K*_D_ value of 0.2 µM
- A generalizable TR-FRET platform enables multimodal profiling of BTB proteins
- The investigational drug CDDO is a partial antagonist of the KEAP1-CUL3 interaction

## Introduction

The Kelch-like family of E3 ubiquitin ligase adaptor proteins (KLHL1-42) comprising BTB, BACK and Kelch domains are associated with a wide range of chronic diseases, including autoimmune and inflammatory diseases, neurodegeneration and cancer [1-3]. Most studied as a therapeutic target is the protein KEAP1 (KLHL19), which regulates the anti-oxidant response by promoting the ubiquitination and proteasomal degradation of substrates, including the NRF2 transcription factor [4-7]. Oxidative and electrophilic stress induce cysteine modifications that disrupt KEAP1 function, allowing NRF2 to accumulate for the transactivation of stress-response genes [8-10].

Ubiquitination by KEAP1 and other KLHL-family proteins is dependent on their binding to the N-terminal domain of CUL3 (CUL3_NTD_), which acts as a scaffold for their assembly into multisubunit Cullin-RING E3 ligases [11-13]. Following activation by an E1 enzyme, charged E2-ubiquitin conjugates are recruited to the E3 complex by the RING-domain containing RBX1 subunit, which assembles with the CUL3 C-terminal domain (CUL3_CTD_) [14, 15]. Neddylation of the CUL3_CTD_ is predicted to induce conformational changes in the complex that position the ubiquitin moiety optimally for its conjugation to the KEAP1-bound substrate [16, 17].

The structural basis for substrate recruitment by the Kelch domain of KEAP1 has been revealed by numerous co-crystal structures, including structures with both the ‘ETGE’ and ‘DLG’ degron motifs of NRF2 [18-25]. However, to date there are neither structures elucidating the critical interaction between KEAP1 and CUL3, nor data to show the contributions of different domains to their binding affinity. The molecular binding model (see schematic Fig. 1) is therefore currently inferred from the structures of other CUL3_NTD_ complexes, including those of KLHL3 [26], KLHL11 [27], SPOP [28, 29] and the vaccinia virus protein A55 [30], as well as the structure of the isolated BTB domain of KEAP1 [31]. Collectively, the structures identify a common interaction between the BTB domain and 3-box of the E3 and the first Cullin repeat domain of CUL3_NTD_. The 3-box forms a short helical motif that was found to be critical for high-affinity CUL3 interaction (analogous to the F-box and SOCS-boxes in other cullin-based E3s) [29]. The structure of the KLHL11-CUL3 complex showed the 3-box packing at the junction between the BTB and BACK domains and forming a hydrophobic groove that accommodated an N-terminal extension in CUL3 (‘N22’ in Fig. 1) [27]. Notably, deletion of the N-terminal extension resulted in a 30-fold lower affinity, highlighting its importance for the interaction [27].

**Fig. 1.**
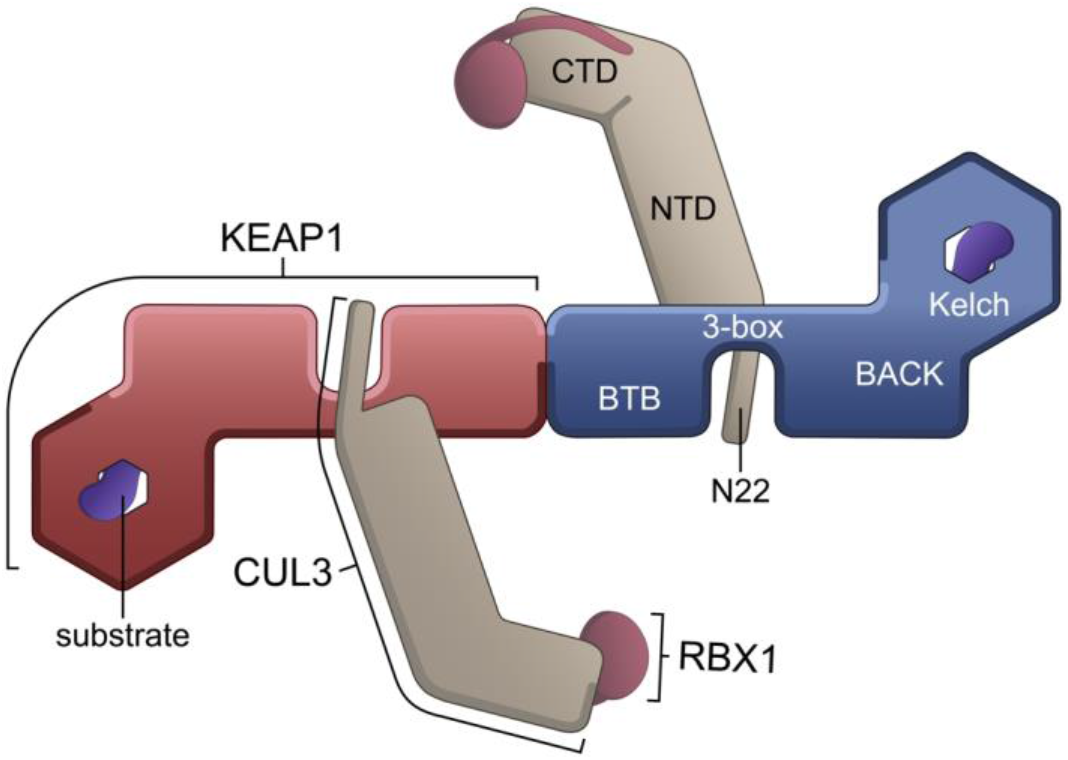
Schematic showing the domain architecture of KEAP1 and CUL3 and predicted model for their interaction. KEAP1 is shown as a homodimer in red and blue. Substrate degrons (purple) are shown bound to the Kelch domains. The CUL3 (light brown) C-terminal domain (CTD) is bound to RBX1 (claret). The CUL3 N-terminal domain (NTD) and 22 residue N-terminal extension (N22) are predicted to bind to the KEAP1 BTB and 3-box domains (model based on the previously determined structure of a KLHL11-CUL3 complex [27]). The BTB domain was first identified as a conserved motif in the Drosophila proteins bric-à-brac, tramtrack and broad complex (reviewed in [58]). Likewise, the Kelch repeat domain was first identified in the Drosophila Kelch protein (reviewed in [59]). The BACK domain (for BTB and C-terminal Kelch) is also known as the intervening region (IVR) in KEAP1 and includes the 3-box motif at it N-terminus [60].

Plant-derived triterpenoid drugs, including 2-cyano-3,12-dioxooleana-1,9(11)-dien-28-oic acid (CDDO, bardoxolone), and its methyl ester CDDO-Me (bardoxolone methyl), have been postulated to restrict the interaction of KEAP1 with CUL3 and thereby stabilize NRF2 for cytoprotection [31, 32]. This has led to the clinical investigation of CDDO-Me in conditions such as cancer, chronic kidney disease, pulmonary hypertension and COVID-19 [5, 33-36]. A co-crystal structure of CDDO revealed its covalent binding to Cys151 in a shallow pocket in the KEAP1 BTB domain [31], which we subsequently showed to bind reversibly with a *K*_D_ value of 3 nM [37]. However, from the crystallographic data, it remained unclear whether CDDO acts to restrict CUL3 binding – possibly via steric hindrance or via induced conformational changes [31]. Of note, a study utilizing fluorescence recovery after photobleaching (FRAP) in live cells did not establish support that CDDO abolishes KEAP1-CUL3 interaction, raising further uncertainty about the proposed mechanism of action [38]. There have also been conflicting reports on the stoichiometry of the KEAP1-CUL3 complex [39, 40]. As the BTB domain of KEAP1 forms a homodimer, it is expected that the homodimer will afford binding sites for two CUL3 proteins (Fig. 1) [27, 39]. However, at least one study has suggested that only one CUL3 protein is bound [40].

In this study, we aimed to provide a structural model of the KEAP1-CUL3 complex and to establish a robust assay system to measure their interaction affinity and the effect of CDDO.

We determined a crystal structure of the BTB and 3-box domains of KEAP1 in complex with the CUL3_NTD_ that revealed a heterotetrameric complex with a 2:2 stoichiometry. To support the structural data, we developed a generalizable TR-FRET-based assay system to profile the binding of BTB-domain-containing proteins to CUL3 and determine the contribution of distinct protein features, revealing the importance of the CUL3 N-terminal extension for high affinity binding. We further provide direct evidence that CDDO does not disrupt the KEAP1-CUL3 complex, even at high concentrations, but rather reduces the affinity of the KEAP1-CUL3 interaction.

## Results

### Structure determination

To determine the structural mechanisms of the KEAP1-CUL3 interaction, we prepared recombinant proteins with various truncations to identify regions compatible with crystallization. Crystals were obtained following purification of a complex consisting of the BTB and 3-box regions of human KEAP1 (residues 48-213; herein KEAP1_BTB-3-box_) and the N-terminal domain of CUL3 (residues 1-388; herein CUL3_NTD_), consistent with the expected interaction domains (Fig. 1). Larger complexes comprising all folded domains of KEAP1 (residues 48-624, herein KEAP1_BTB-BACK-Kelch_) or the full-length CUL3-RBX1 complex did not yield crystals.

The resulting structure was solved by molecular replacement in space group C2 2 2_1_ and refined at 3.45 Å resolution (see Table S1 for data collection and refinement statistics). A single chain each of KEAP1 and CUL3 was identified in the crystallographic asymmetric unit. The electron density maps allowed KEAP1 to be modelled from residues 51-204 and the CUL3 chain from residues 26-381, except for a disordered loop between residues 331 and 338. Crystallographic symmetry revealed the expected homodimerization of the KEAP1 BTB domain yielding an overall KEAP1-CUL3 heterotetrameric complex with a 2:2 stoichiometry and overall complex dimensions of 162 × 90 × 43 Å (Fig. 2).

**Fig. 2.**
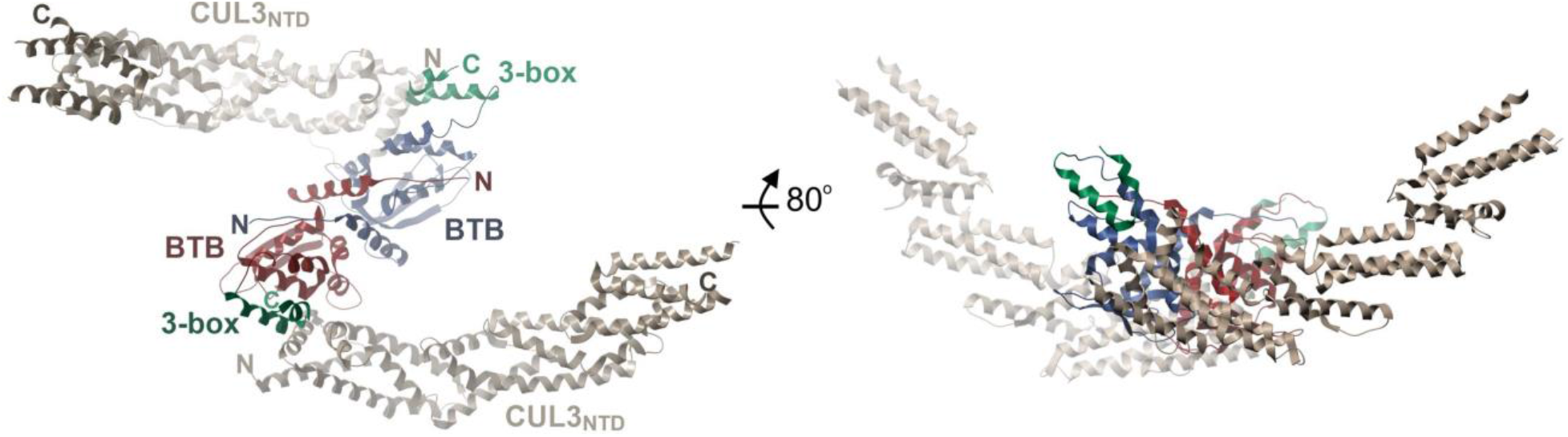
Structure of the crystallised KEAP1-CUL3 complex. The structure shown in two orientations reveals a stoichiometry of 2:2 for KEAP1_BTB-3-box_ binding to the Cul3_NTD_.

### Interactions in the KEAP1-CUL3 interface

The KEAP1 BTB and 3-box domains were bound exclusively to the first Cullin repeat domain of CUL3 (Fig. 3A). Compared to the free BTB structure [31], the KEAP1 BTB domain exhibited an induced fit characterized by alternative packing of the α3-β4 loop to insert KEAP1 Leu115 into a deep hydrophobic pocket formed between the H2 and H4 helices of CUL3 (Fig. 3A). Of note, Leu115 showed the highest buried interface area of any residue in the complex (Fig. 3B-C) and was displaced by 9 Å compared to the free KEAP1 structure (Fig. 3A). Leu115 belongs to a φ-x-E motif first defined in the SPOP-CUL3 structure, and conserved in BTB family E3 ligases [26, 28], where Leu115 represents the hydrophobic residue φ, Arg116 is the charged/polar residue x and Glu117 is the conserved glutamate forming hydrogen bond interactions with the CUL3 H2 helix. The H2 and H5 helices of CUL3 are also notable for a cluster of tyrosine residues that form hydrogen bonds with the KEAP1 α5 and α7 helices in the BTB and 3-box domains, respectively (Fig. 3A). Superposition of the structure with the KEAP1_BTB_-CDDO complex revealed that the CDDO binding site was on the opposite face of the BTB domain to the CUL3 interface and therefore was unlikely to directly disrupt these observed interactions (Fig. 3A).

**Fig. 3.**
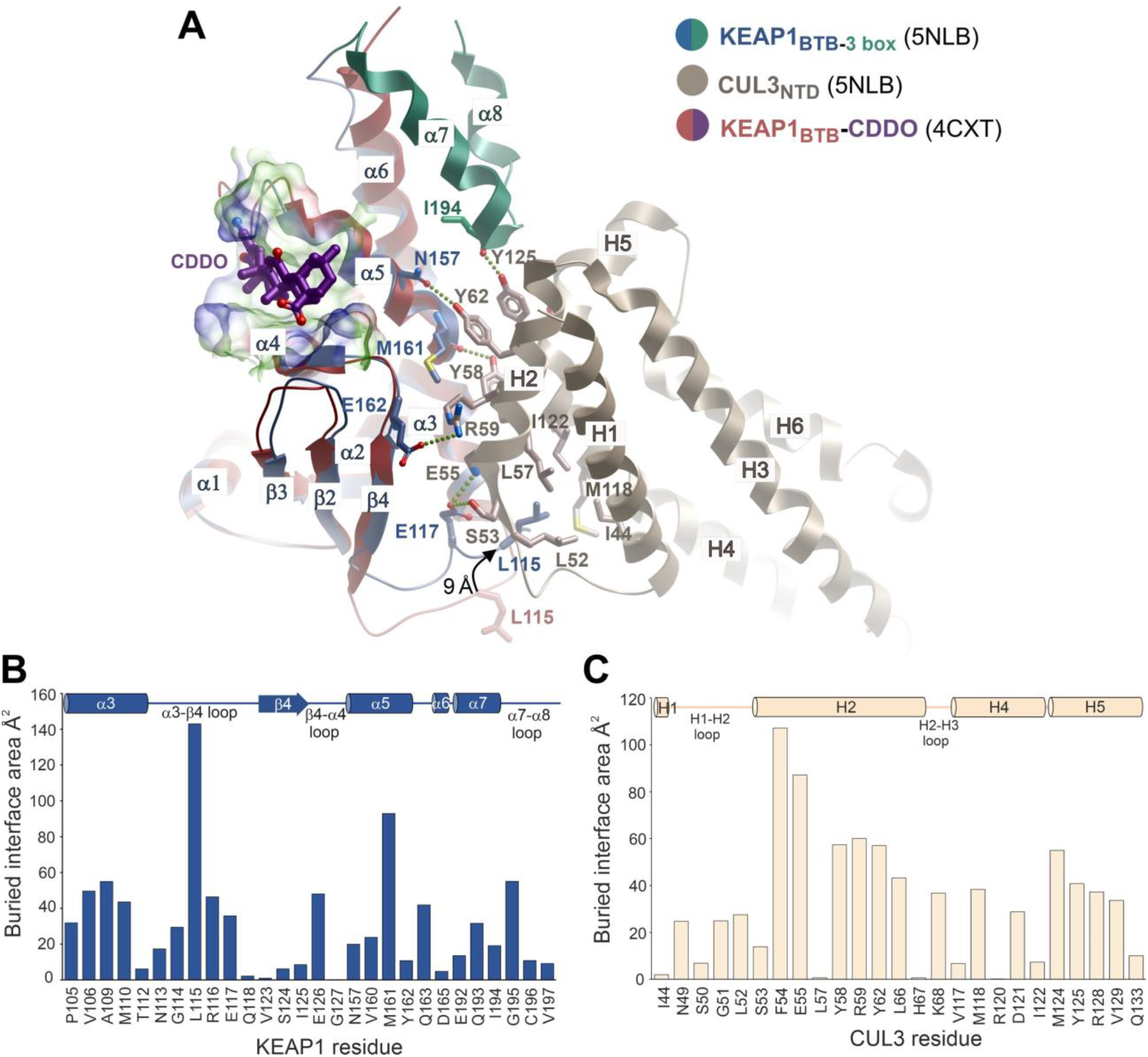
Interactions in the KEAP1-CUL3 interface. (A) Superposition of the KEAP1-CUL3 complex with the previously determined structure of the KEAP1 BTB domain bound to CDDO. Different structural features are coloured according to the key. An arrow highlights conformational differences in the BTB α3-β4 loop in the two structures as highlighted by the 9Å difference in the position of KEAP1 Leu115. Selected hydrogen bond (dotted lines) and hydrophobic interactions are shown in the KEAP1-CUL3 complex. A surface representation of the CDDO-binding pocket is shown with partial transparency. CDDO is shown as purple sticks. (B) Buried interface areas of KEAP1 residues bound to CUL3 calculated using the Protein interfaces, surfaces and assemblies server (PISA) at the European Bioinformatics Institute (http://www.ebi.ac.uk/pdbe/prot_int/pistart.html)[61]. Residue numbers and secondary structure elements are indicated. (C) Buried interface areas of CUL3 residues bound to KEAP1.

### Comparison with the extended interface of the KLHL11-CUL3 structure

Overall, the KEAP1-CUL3 interface buried a surface area of 833 Å^2^. By comparison, the structure of a KLHL11_BTB-BACK_ construct bound to CUL3_NTD_ showed an extended interface of 1508 Å^2^ boosted by additional interactions between an N-terminal CUL3 extension (“N22” in Fig. 1) and a hydrophobic groove formed between KLHL11 α5 (BTB) and α7 (3-box) (Fig. 4A-B) [27]. No electron density was observed for the same CUL3 N-terminal region in the KEAP1 complex despite the sequence similarity of the binding site (Fig. 4A). This might reflect hindrance from crystal packing (Fig. S1), or the absence of the full BACK domain in the crystallized KEAP1 construct, which is likely to stabilze the 3-box structure and form further minor contact with CUL3. In addition, the KEAP1 3-box contains some bulkier substitutions, such as Phe190, that could diminish the size of the hydrophobic groove for interaction (Fig. 4B). While the binding of the CUL3 N-terminal extension was not observed in the KEAP1 complex structure, modelling of this region using the equivalent KLHL11_BTB-BACK_ co-structure suggested a potential steric clash between the CUL3 extension and the small molecule inhibitor CDDO (Fig. 4B), providing one possible explanation for its mode of action.

**Fig. 4.**
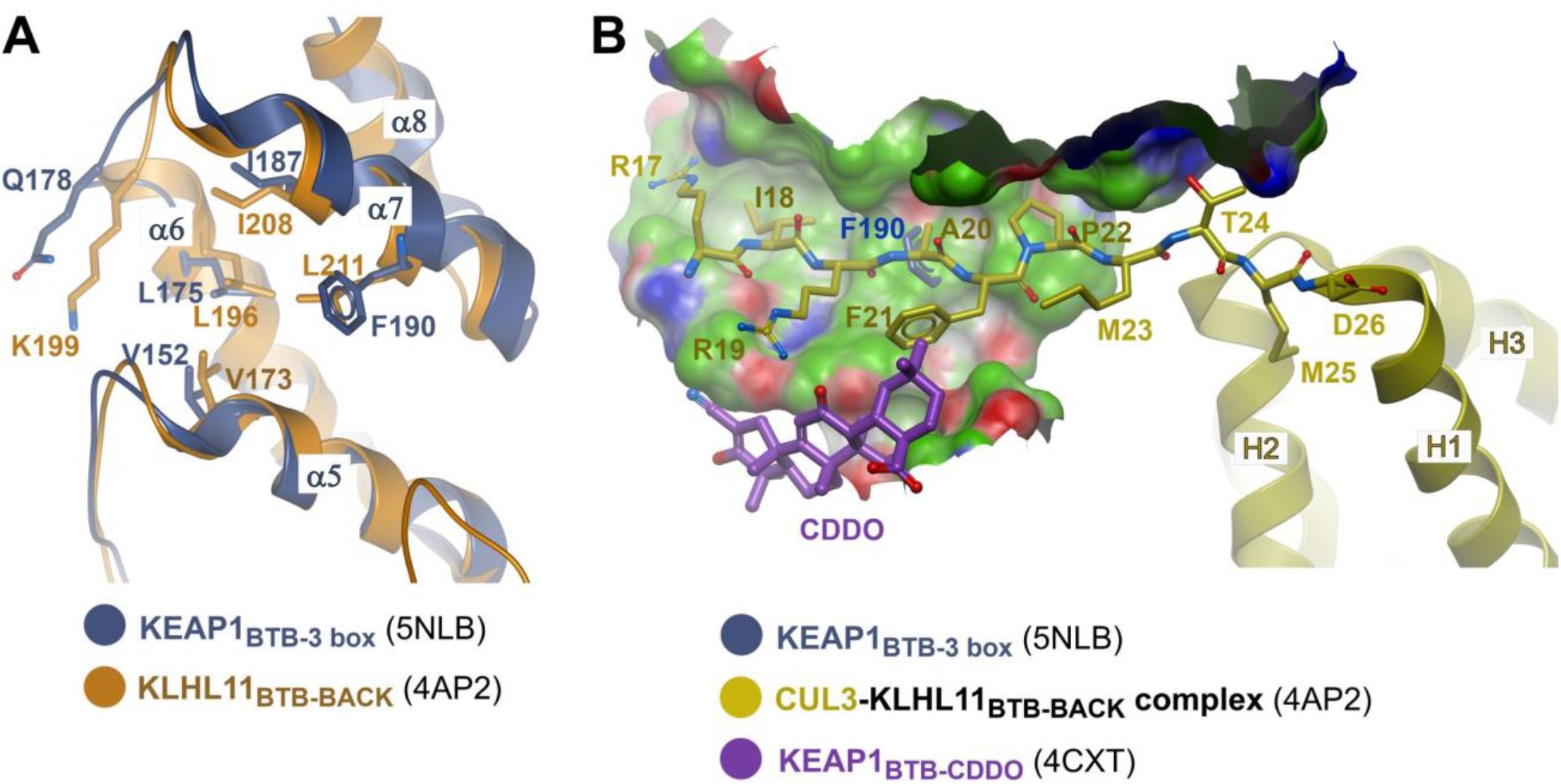
Comparison of the 3-box hydrophobic groove in the respective KEAP1 and KLHL11 bound CUL3 complexes. (A) Superposition of KEAP1 (dark blue) and KLHL11 (orange) showing conservation of a hydrophobic groove between the BTB domain (α5) and 3 box (α7). KLHL11 Leu211 is replaced by Phe190 in KEAP1. (B) Superposition of CUL3 complexes of KEAP1 and KLHL11 with the KEAP1-CDDO complex. A surface representation of the 3-box hydrophobic groove in the KLHL11-CUL3 complex is shown (green with blue and red areas denoting hydrogen bond donor and acceptor positions, respectively) with interacting CUL3 residues shown as yellow sticks. Superposition predicts clashes between KEAP1 Phe190 (protruding through this surface) and CUL3 Ala20 as well as between KEAP1-bound CDDO and CUL3 Phe21.

### Biolayer interferometry indicates that KEAP1_BTB-3-box_ binds to CUL3 relatively weakly

Next, we aimed to determine the binding affinity between KEAP1 and CUL3, and to evaluate the contributions of the CUL3 N-terminal extension, as well as the BTB-BACK and Kelch domains of KEAP1. We first used biolayer interferometry (BLI) to profile the interaction of the proteins used in the structure determination. Biotinylated CUL3_NTD_ was captured on a streptavidin-functionalized sensor and binding was quantified using serial dilutions of KEAP1_BTB-3-box_. The apparent *K*_D_ = 1.7 μM (95% CI 1.0-2.5 μM; Fig. 5) determined under steady-state conditions was comparable to the results obtained with a reverse setup immobilizing biotinylated KEAP1_BTB-3-box_ and titrating serial dilutions of the CUL3_NTD_ protein (Fig. S2A), or the full length CUL3-RBX1 complex (Fig. S2B). The measured binding affinity was markedly weaker than those reported for CUL3_NTD_ binding to KLHL11_BTB-BACK_ (*K*_D_ = 20 nM [27]) or to a SPOP_BTB-3-box_ construct (*K*_D_ = 17 nM [29]). Instead, the measured interaction was similar to previous studies using CUL3_NTD_ constructs with an N-terminal deletion (either CUL3_NTDΔN19_ or CUL3_NTDΔN22_), including those analyzing binding to the SPOP BTB domain alone (*K*_D_ = 1.0 μM [28]), or to a lesser extent KLHL11_BTB-BACK_ (*K*_D_ = 0.65 μM [27]) or KLHL3_BTB-BACK_ (*K*_D_ = 0.11 μM [26]).

**Fig. 5.**
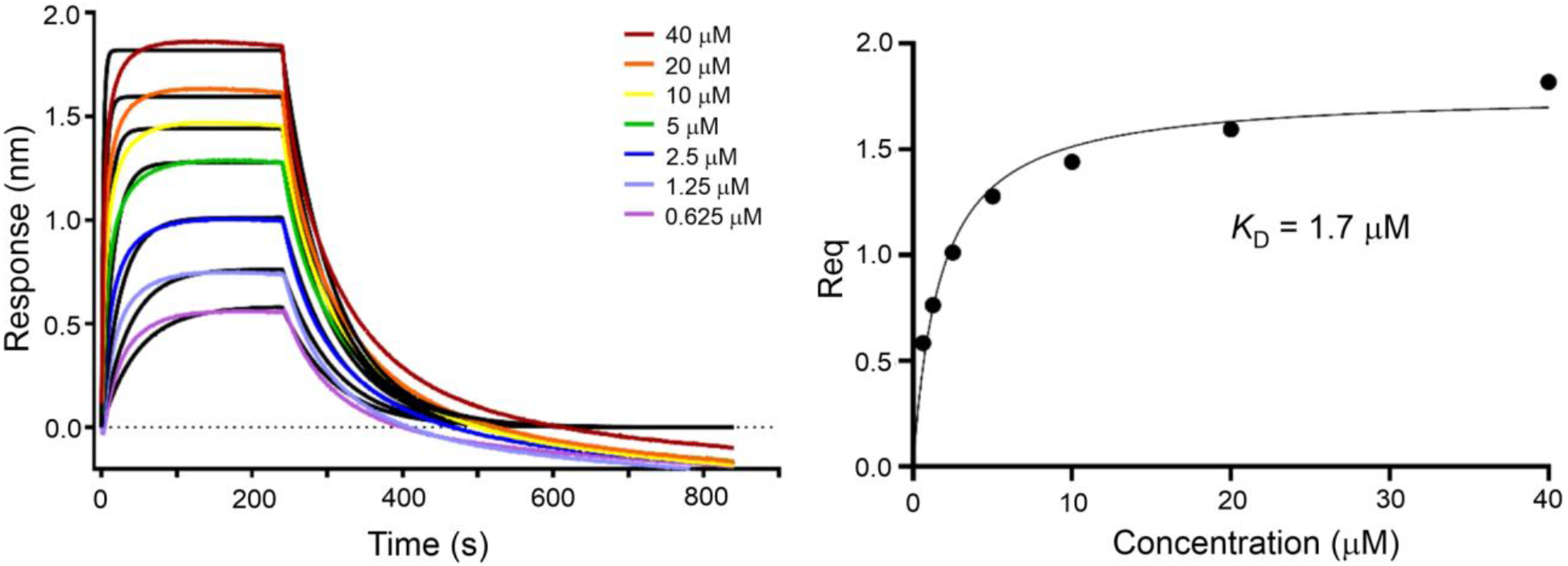
Biolayer interferometry (BLI) measurements of CUL3 binding to the KEAP1_BTB-3-box_. Binding equilibrium and kinetic measurements were determined on an Octet RED384 instrument (FortéBio). Biotinylated CUL3_NTD_ was immobilized on a streptavidin-functionalized sensor tip and binding to serial dilutions of KEAP1_BTB-3-box_ was quantified by fitting to a Langmuirian 1:1 model. Steady state equilibrium analysis yielded an apparent *K*_D_ = 1.7 μM (95% CI 1.0-2.5 μM) and binding kinetics of *k*_on_ = 1.02 × 10^4^ M^-1^s^-1^, *k*_off_ = 1.45 × 10^−2^ s^-1, App^*K*_D_ = 1.4 μM.

Together, these data highlight the importance of the interaction between the CUL3 N-terminal extension and the 3-box grove in BTB-containing proteins such as KLHL11 and SPOP. By extension, the lack of electron density for the CUL3 N-terminus in our co-structure with the KEAP1_BTB-3-box_ may suggest the absence of this binding feature in KEAP1, providing a rationale for the comparatively low binding affinity measured with the KEAP1 constructs. To rule out that these results are not the consequences of the absence of a complete BACK domain, we set out to include KEAP1_BTB-BACK-Kelch_ and an N-terminally truncated CUL3 that lacks all 3-box interacting residues (CUL3_NTDΔN22_) in our analysis. Unfortunately, the KEAP1_BTB-BACK-Kelch_ construct exhibited poor behavior in BLI experiments, precluding characterization of its binding to CUL3_NTD_ (Fig. S2C). Similarly, we were unsuccessful in establishing a functional BLI assay for measuring the binding of KEAP1_BTB-3-box_ to the CUL3_NTDΔN22_ construct (Fig. S2D). We, therefore, explored a TR-FRET-based experimental design as an alternative strategy, following our previous work utilizing CoraFluor-1 as the luminescent donor for the characterization of KEAP1 ligands and KEAP1 homodimerization [37]. Homogenous TR-FRET assays offer several advantages over other biophysical methods and even enable the quantitative measurement of low-affinity interactions.

### TR-FRET assays reveal the importance of other KEAP1 domains for CUL3 interaction

We rationalized that pairwise labeling of BTB-containing proteins and CUL3_NTD_ would provide a straightforward and target-agnostic strategy for the characterization of binding affinities (Fig. 6A-D). TR-FRET donor and acceptor functionalization was accomplished through direct acylation using CoraFluor-1-Pfp and AF488-Tfp, respectively [41-43]. We selected direct chemical labeling over the use of labeled anti-epitope tag antibodies or streptavidin, which can complicate data interpretation due to the formation of higher-order complexes. To validate our approach, we first performed a saturation binding experiment with CoraFluor-1-labeled KLHL11_BTB-BACK_ and AF488-CUL3_NTD_, which yielded a *K*_D_ value of 20 nM (95% CI 19-22 nM), consistent with our previous ITC data (Fig. 6B) [27]. Direct measurement of the dissociation rate constant (*k*_off_ = 3.81 × 10^−3^ s^-1^) was determined by the addition of an excess of the respective unlabeled competitor to preequilibrated TR-FRET donor and acceptor functionalized protein complexes, establishing an assay equilibration time of ∼15 min (5 × *t*_1/2_) (Fig. 6C) [44, 45]. Dose-response titration of unlabeled CUL3_NTD_ or KLHL11_BTB-BACK_ as competitors yielded similar *K*_D_ values (Fig. 6D) and provided evidence that dye functionalization was well tolerated and did not alter the binding affinity (Table S2).

**Fig. 6.**
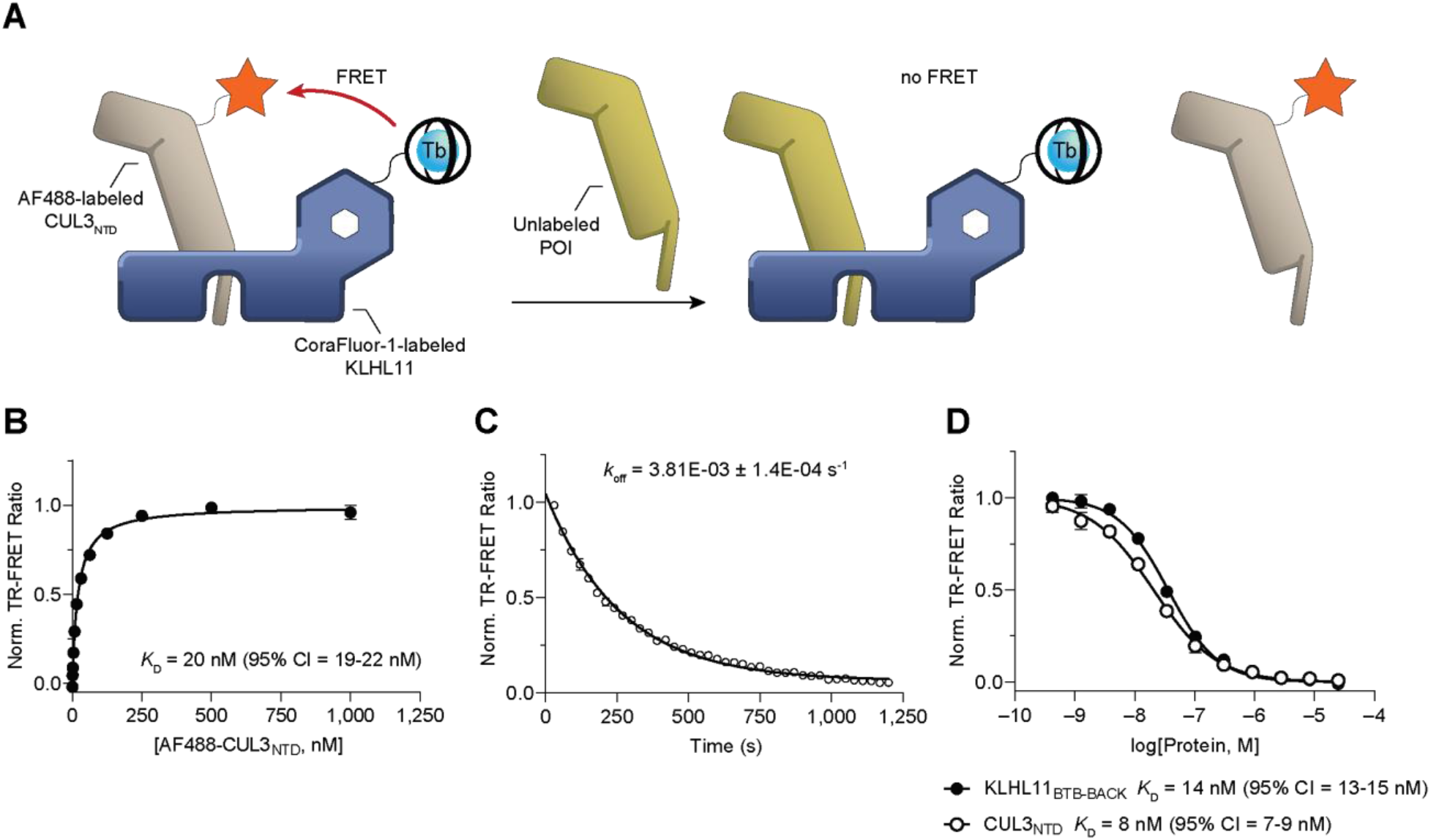
Validation of TR-FRET assay approach with KLHL11_BTB-BACK_ and CUL3_NTD_. (A) Schematic of the TR-FRET assay principle with CoraFluor-1-labeled KLHL11 and AF488-labeled CUL3_NTD_. Addition of unlabeled protein of interest (POI) competes with either CUL3_NTD_ (shown) or KLHL11 (not shown), disrupting the TR-FRET process. For clarity, only one KLHL11_BTB-BACK-Kelch_ subunit from the KLHL11 homodimer is shown. (B) Saturation binding of AF488-labeled CUL3_NTD_ to CoraFluor-1-labeled KLHL11_BTB-BACK_ yielded a *K*_D_ value of 20 nM, consistent with our previous ITC data [27]. (C) Determination of the dissociation rate constant (*k*_off_ = 3.81×10^−3^ s^-1^) establishes an assay equilibration time of ∼15 min (5 × *t*_1/2_). (D) Dose-response titration of unmodified KLHL11_BTB-BACK_ and CUL3_NTD_ in TR-FRET protein displacement assay with AF488-labeled CUL3_NTD_ and CoraFluor-1-labeled KLHL11_BTB-BACK_.

Because KEAP1 and KLHL11 bind to the same site of CUL3_NTD_, this assay system is also suitable as a ligand displacement assay for profiling the binding affinity of KEAP1 constructs and, by extension, other BTB domain-containing proteins. However, some BTB proteins have previously been reported to form heterodimers, which could result in a non-linear response of this assay system and potentially misleading data [3]. Therefore, we first employed our previously reported KEAP1 dimerization assay to address this question and test the capacity of KLHL11 to form heterodimers with KEAP1 [37]. As shown in Fig. 7A, we did not observe KEAP1-KLHL11 heterodimerization, rendering our approach viable for the direct profiling of KEAP1-CUL3_NTD_ interaction with this assay platform.

**Fig. 7.**
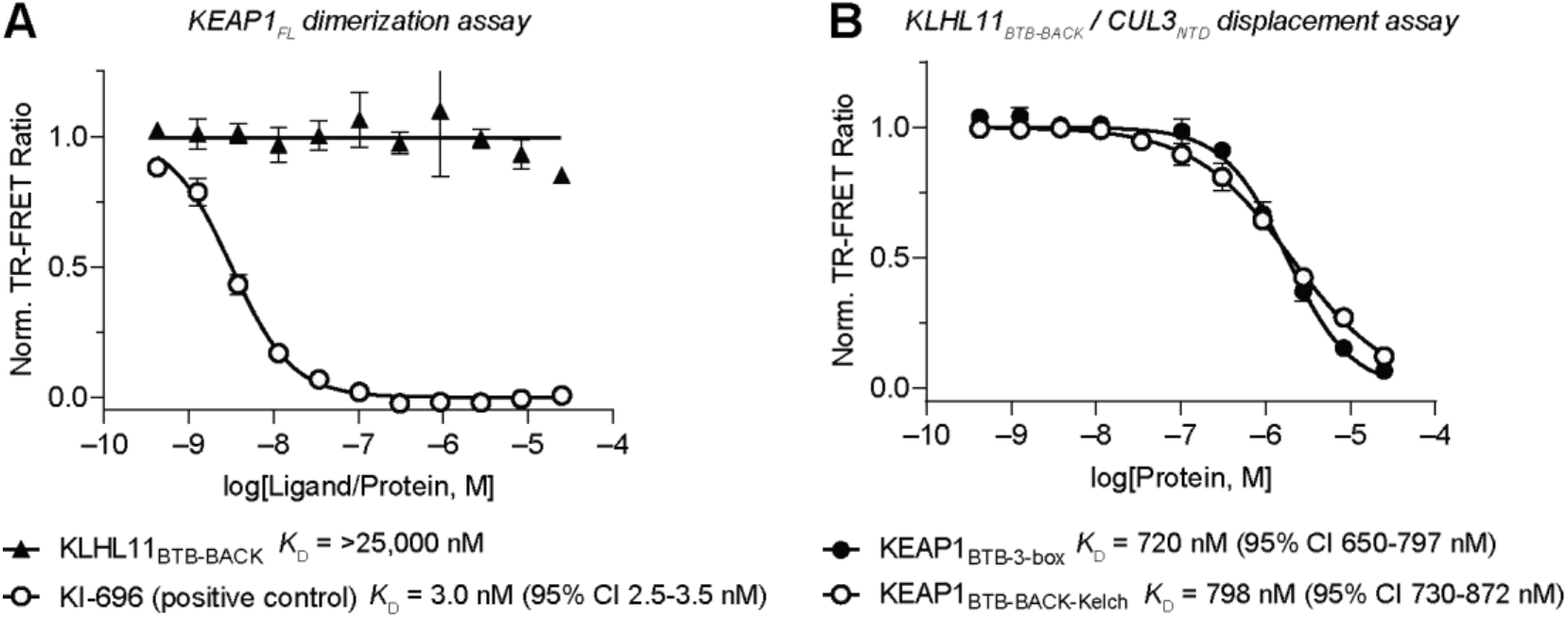
A generalizable TR-FRET platform for profiling BTB domain-containing proteins. (A) Dose-response titration of KLHL11_BTB-BACK_ and small molecule KEAP1-Kelch domain inhibitor KI-696 (positive control) in KEAP1_FL_ dimerization assay [37, 62], using a mixture of CoraFluor-1- and FITC-labeled Nrf2-derived peptides (LDEETGEFL-CONH_2_), showing that KLHL11_BTB-BACK_ does not heterodimerize with KEAP1_FL_ at concentrations up to 25 µM. (B) Dose-response titration of KEAP1_BTB-3-box_ and KEAP1_BTB-BACK-Kelch_ in KLHL11_BTB-BACK_-CUL3_NTD_ TR-FRET protein displacement assay provides evidence that the complete BTB-BACK domain does not meaningfully contribute to the binding affinity of the KEAP1-CUL3_NTD_ interaction.

Next, we performed dose-response experiments with the respective KEAP1 constructs in the KLHL11-CUL3_NTD_ protein displacement assay. We determined *K*_D_ values of 720 nM (95% CI 650-797 nM) and 798 nM (95% CI 730-882 nM) for KEAP1_BTB-3-box_ and KEAP1_BTB-BACK-Kelch_, respectively, suggesting that the complete BTB-BACK domain does not meaningfully contribute to the binding affinity of the KEAP1-CUL3_NTD_ interaction compared to BTB-3-box alone (Fig. 7B). Notably, these results provide an absolute comparison of the respective binding affinities because the titrations of unlabeled KLHL11 and KEAP1 are performed with the same reporter system.

Following the validation of our assay approach, we developed a similar TR-FRET protein displacement assay using full-length, untagged KEAP1 (KEAP1_FL_ residues 1-624) and CUL3_NTD_ to establish a complete characterization of the KEAP1-CUL3_NTD_ interaction. Saturation binding of CoraFluor-1-labeled KEAP1_FL_ and AF488-labeled CUL3_NTD_ yielded a *K*_D_ value of 222 nM (95% CI 211-234 nM; Fig. 8A), suggesting that the N-terminal 48 amino acids of KEAP1 may contribute to higher affinity KEAP1-CUL3_NTD_ binding. Kinetic analysis provided a *k*_off_-value of 5.04 × 10^−3^ s^-1^ (assay equilibration time ∼12 min; Fig. 8B), and dose-response titration of KEAP1_BTB-3-box_ and KEAP1_BTB-BACK-Kelch_ yielded similar affinity values as measured in our orthogonal assay (*K*_D_ = 1,042 nM [95% CI 943-1151 nM] and 396 nM [95% CI 363-430 nM], respectively) (Fig. 8C, Table S3).

**Fig. 8.**
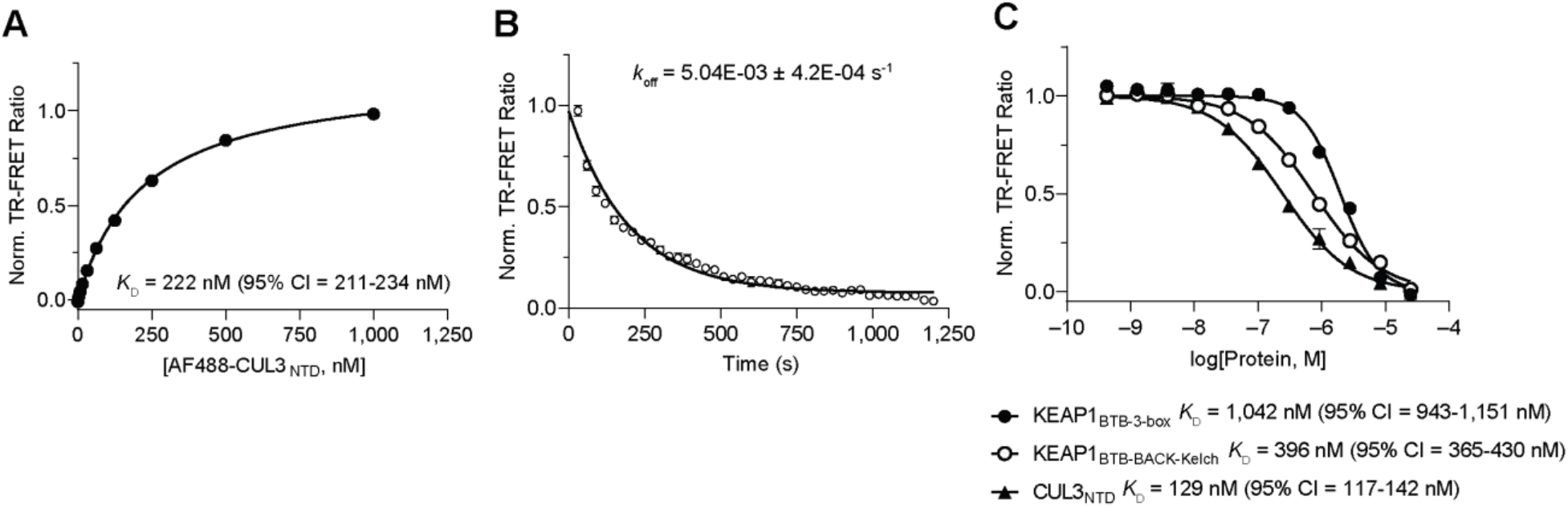
KEAP1_FL_-CUL3_NTD_ TR-FRET protein displacement assay further characterizes KEAP1-CUL3NTD interaction. (A) Saturation binding of AF488-labeled CUL3_NTD_ to CoraFluor-1-labeled KEAP1_FL_ yielded a *K*_D_ value of 222 nM, suggesting that the N-terminal 48 residues of KEAP1_FL_ contribute to slightly higher binding affinity to CUL3_NTD_. (B) *k*_off_ determination (koff = 5.04×10^−3^ s^-1^) establishes an assay equilibration time of ∼12 min (5 × *t*_1/2_). (C) Dose-response titration of unmodified KEAP1_BTB-3-box_, KEAP1_BTB-BACK-Kelch_, and CUL3_NTD_ in TR-FRET protein displacement assay with AF488-labeled CUL3_NTD_ and CoraFluor-1-labeled KEAP1_FL_.

To assess the relevance of the N-terminal extension in CUL3, we employed our suite of TR-FRET protein displacement assays to characterize the affinity of CUL3_NTDΔ22_ for both KEAP1_FL_ and KLHL11_BTB-BACK_ (Fig. 9). We found that the lack of the N-terminus in CUL3 decreased the affinity for KLHL11_BTB-BACK_ by > 200-fold (*K*_D_ = 1,840 nM [95% CI 1,702-1,990 nM]), which is even more pronounced than the 30-fold reduction estimated by ITC in our previous report [27]. Similarly, we observed a ∼100-fold decreased affinity of KEAP1_FL_ for CUL3_NTDΔ22_ (*K*_D_ = 13,783 nM [95% CI 12,047-16,119 nM]) compared with CUL3_NTD_ (*K*_D_ = 129 nM [95% CI 117-142 nM]). This result was unexpected because of the lack of electron density for the 22 amino acid N-terminal extension in our KEAP1_BTB-3-box_-CUL3_NTD_ co-crystal structure. Nonetheless, our data support the role of the N-terminus in mediating high-affinity interactions between the KEAP1/KLHL11 3-box grooves and CUL3, and suggest that the complete BACK domain might be required for structured binding.

**Fig. 9.**
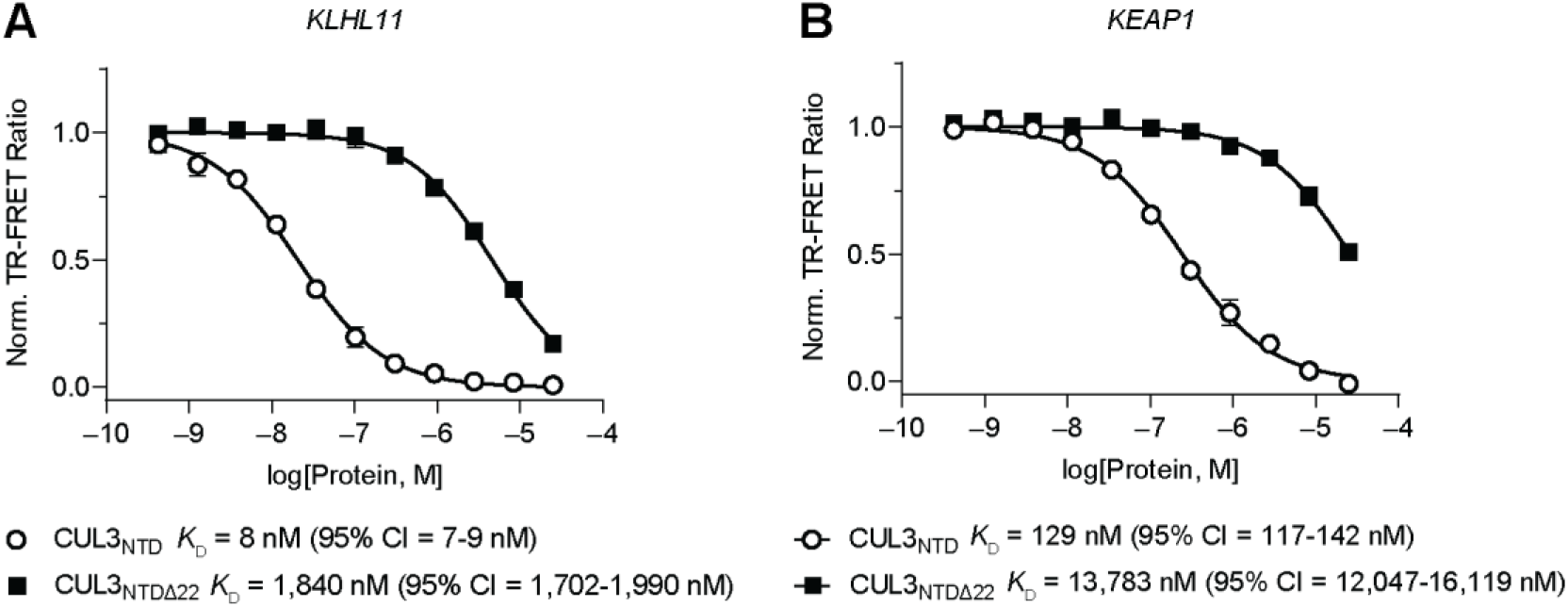
Examining the role of the CUL3_NTD_ N-terminal extension on binding affinity toward KLHL11 and KEAP1. (A-B) Dose-response titration of CUL3_NTD_ and CUL3_NTDΔ22_ in TR-FRET protein displacement assay between AF488-labeled CUL3_NTD_ and (A) CoraFluor-1-labeled KLHL11_BTB-BACK_, or (B) CoraFluor-1-labeled KEAP1_FL_. Deletion of the N-terminal 22 amino acid residues results in a ∼100-fold loss of binding affinity of CUL3_NTD_ toward both proteins.

### TR-FRET reveals partial inhibition by CDDO

We have previously shown that CDDO binds to the KEAP1 BTB with *K*_D_ = 3 nM but does not interfere with KEAP1 dimerization [37]. To understand if CDDO can inhibit KEAP1-CUL3 complex formation, we used our TR-FRET platform and evaluated CDDO in dose-response. We found a dose-dependent decrease in signal intensity that plateaued at 50% “inhibition” at high compound concentrations (Fig. 10A). Although this type of “incomplete” inhibition can be observed for small molecules that are insufficiently soluble at concentrations above the IC_50_ in the respective assay system, this is not the case for CDDO. An alternative explanation for this observation is that CDDO binding alters the affinity between KEAP1 and CUL3. We, therefore, determined the binding affinity of KEAP1_FL_ to CUL3_NTD_ in the absence and presence of high concentrations of CDDO. The saturation binding experiments revealed that CDDO decreases KEAP1-CUL3_NTD_ binding affinity by > 2-fold, functioning only as a partial antagonist that cannot completely disrupt the complex (Fig. 10B). Although this shift in affinity might be sufficient to modulate the function of this critical redox sensor in cells to trigger activation of the ARE (antioxidant response element), it does not rule out more pronounced functional consequences as the result of a distorted orientation of CUL3 with respect to the protein substrate.

**Fig. 10.**
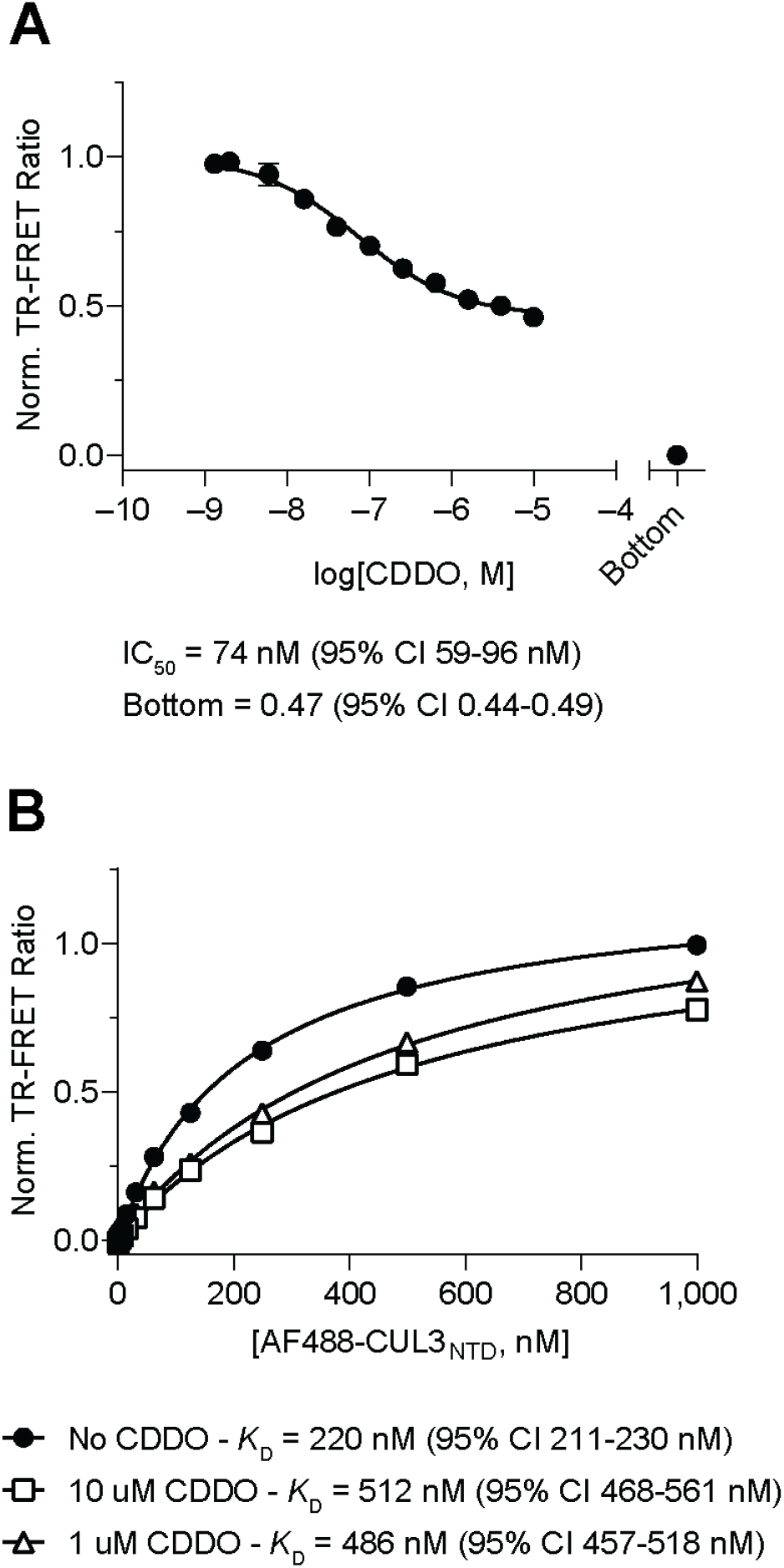
CDDO is a partial antagonist of the KEAP1-CUL3 interaction. (A) Dose-response titration of CDDO in protein displacement assay with AF488-labeled CUL3_NTD_ and CoraFluor-1-labeled KEAP1_FL_ yielded an IC_50_ value of 74 nM and incomplete inhibition. Assay floor determined with 25 µM CUL3_NTD_. (B) Saturation binding of AF488-labeled CUL3_NTD_ to CoraFluor-1-labeled KEAP1_FL_ in the absence or presence of 1 or 10 µM CDDO reveals that CUL3_NTD_ has >2-fold reduced affinity toward the KEAP1-CDDO binary complex compared to KEAP1 alone.

## Discussion

Cullin-RING ligase complexes typically utilize separate protein subunits as Cullin adaptor and substrate receptor. For example, the SCF (SKP1-CUL1-F-box) class use SKP1 as an adaptor to link CUL1 with an F-box-containing protein that functions as the substrate receptor [14]. CUL3 complexes form an exception in which both functionalities are incorporated into a single protein (e.g. KEAP1), allowing greater sequence diversity in their Cullin interfaces [27, 46]. While the sequences of the KEAP1 BTB and 3-box domains have diverged from other BTB-containing proteins (e.g. 26.5% sequence identity across these domains in KLHL11), their structure in complex with CUL3_NTD_ shows a conserved mechanism of assembly, including a heterotetrameric packing arrangement with 2:2 stoichiometry and an induced fit of the KEAP1 α3-β4 loop.

Except for KLHL11, all previously determined co-structures used truncated forms of CUL3 lacking the N-terminal extension. Thus, in our new structure of the KEAP1-CUL3 complex, we expected to observe the packing of the N-terminal CUL3 region (‘N22’), similar to the KLHL11 co-structure, that would provide a better understanding of the predicted interaction with the 3-box groove and establish a molecular basis for the antagonistic effects of CDDO. However, this region appears to be disordered in the new structure, with the first defined CUL3 residue (Asp26) located some 19 Å from the CDDO binding site, where its interaction with KEAP1 is unlikely to be affected. The 3-box groove appears to be slightly shallower in KEAP1 than in KLHL11. Cleasby et al. previously postulated that the CUL3 N-terminal extension might instead bind to the KEAP1 Cys151 site and therefore compete more directly with CDDO [31]. However, our structure provides no evidence to support this prediction. Nonetheless, in agreement with Cleasby et al., we observe a potential steric clash with CUL3 when modelling its binding to the 3-box groove. To gain insights into the contributions of different domains and the impact of CDDO on KEAP1-CUL3 binding, we determined the equilibrium binding constants and binding kinetics of the various constructs. Unfortunately, we were unable to establish functional biolayer interferometry assays for several protein combinations. However, we successfully developed a robust and versatile biochemical TR-FRET assay platform based on our CoraFluor technology that facilitated the comprehensive, quantitative measurement of the CUL3 interactions. Importantly, when available, the data obtained by TR-FRET were in good agreement with the results determined by BLI.

Although the CUL3 N-terminal extension was not defined in the electron density maps of the KEAP1-CUL3 complex, its deletion still resulted in >100-fold loss of affinity, demonstrating its importance for the interaction. Notably, this affinity differential is comparable to the results obtained for KLHL11-CUL3, for which the CUL3 N-terminal extension is well-refined in the co-complex structure. However, it should be noted that KLHL11 binds significantly (∼10-fold) tighter to the CUL3 constructs than KEAP1. Additionally, we observed a modest contribution to the binding affinity in the presence of the KEAP1 full N-terminus and Kelch domain, as apparent by the comparison of full-length and truncated KEAP1 (*K*_D,full-length_ ∼ 0.2 μM vs. *K*_D,BTB-3-box_ ∼ 1 μM), which might be the consequence of more stable folding of the BTB and 3-box domains within the full length protein.

Furthermore, our TR-FRET assay approach also allowed us to examine the effect of CDDO on KEAP1-CUL3 binding. The KEAP1-CUL3 module has been recognized as a primary target of the cysteine-reactive oleanolic acid derivative CDDO and its analogs. However, the precise mechanism of how CDDO interferes with the function of KEAP1-CUL3 has still not been completely understood. It has previously been shown by crystallography that CDDO binds covalently to Cys151 within the BTB domain of KEAP1 [31]. Based on this and other observations, various modes of action have been proposed, including the disruption of KEAP1 dimer formation and inhibition of KEAP1-CUL3 binding [31, 47-49]. Our recent studies showed that CDDO does not disrupt KEAP1 dimerization [37]. Here, we further demonstrated that CDDO does not disrupt KEAP1-CUL3 binding. Instead, we found that CDDO can act as a partial antagonist that appears to lower the affinity of CUL3 for KEAP1, but does not block the binding of the two proteins. Our findings are consistent with CDDO acting as an allosteric competitive inhibitor that interferes with binding of the CUL3 N-terminal domain to the 3-box groove of KEAP1, which we have shown significantly contributes to the binding affinity of CUL3 and KEAP1. Although we cannot rule out an alternative mechanism. The reversible addition of CDDO or CDDO-me on the thiol of KEAP1 Cys151 has not been detected by mass spectrometry, nor on any other residue within the full length protein [50, 51]. However, irreversible binding of the analog CDDO-epoxide has been mapped to KEAP1 cysteines at positions 257, 273, 288, 434, 489, and 613, both in vitro and in living cells, which could affect multiple protein-protein interactions [50]. Thus, at higher concentrations, it is also possible that CDDO derivatives might be binding to other Cys-side chains of KEAP1 (and/or CUL3), causing structural changes to alter the binding affinity (similar to the absence of the complete N- and C-termini of KEAP1). However, further studies will be needed to explore this activity in greater detail.

Together, the presented structural and biochemical data show the importance of the modular domains of KEAP1 and CUL3 for their heteromeric assembly. KEAP1 represents only one of nearly 200 BTB-containing proteins that can potentially assemble with CUL3 [52]. The established TR-FRET assay system offers a generalizable platform for profiling this protein class and may form a suitable screening platform for ligands that disrupt these interactions by targeting the BTB or 3-box domains to block E3 ligase function.

## Materials and Methods

### Constructs

For bacterial expression, human KEAP1_BTB-3-box_ (Uniprot Q14145, residues 48-213) was cloned into the vector pNIC28-Bsa4, which provides an N-terminal 6xHis tag with a TEV protease cleavage site. A further construct enabling biotinylation of KEAP1_BTB-3-box_ was cloned into the pNIC-Bio3 vector, which provides a TEV-cleavable 6xHis tag and a C-terminal avi tag for biotinylation. Bacterial expression constructs for human KLHL11_BTB-BACK_ (Q9NVR0, residues 67-340 in vector pNIC28-Bsa4), CUL3_NTD_ (Q13618, residues 1-388 in pNIC-CTHF) and CUL3_NTDΔ22_ (residues 23-388 in pNIC-CTHF) were described previously [27]. Vector pNIC-CTHF provides C-terminal 6xHis and FLAG tags cleavable by TEV protease. A construct enabling biotinylation of CUL3_NTD_ (residues 1-389) was cloned into the pNIC-Bio3 vector. All CUL3_NTD_ constructs included mutations I342R and L346D designed to stabilise the isolated N-terminal domain (NTD) as part of the “split-n-express” strategy outlined by Zheng et al. [14].

For baculoviral expression, KEAP1_BTB-BACK-Kelch_ (residues 48-624) and RBX1 (P62877 residues 1-108) were cloned into the baculoviral transfer vector pFB-LIC-Bse providing a TEV-cleavable N-terminal 6xHis tag. Full length CUL3 (residues 1-768) was cloned into pFB-CT6HF-LIC, which provides C-terminal 6xHis and FLAG tags cleavable by TEV protease. Full-length KEAP1 was purchased from Sino Biological (cat# 11981-HNCB).

### Protein expression and purification

Bacterial protein expression was performed in BL21(DE3)-pRARE2 cells. 2 L cultures were grown to mid log phase in LB or 2xTY media with antibiotic selection, then cooled to 18°C and induced with 0.25 mM IPTG. Cells were harvested by centrifugation and resuspended in 40 mL Binding Buffer (50 mM HEPES pH 7.5, 500 mM NaCl, 5 % glycerol, 5 mM imidazole and 0.5 mM TCEP). Lysozyme (1 mg/mL), PEI (1 mL of 5% stock) and protease inhibitors were added before cell lysis by sonication. After centrifugation of the lysate at 50 000 g, the supernatant was filtered (1.2 µm) and incubated for 30 min with 3 mL nickel sepharose resin equilibrated in Binding Buffer. The column was washed with 50-80 mL Wash Buffer (50 mM HEPES pH 7.5, 500 mM NaCl, 5 % glycerol, 30 mM imidazole and 0.5 mM TCEP) and the protein step eluted in 10 mL fractions of Binding Buffer supplemented with 50, 100, 150 and 250 mM imidazole. The fractions were pooled, diluted one third with Binding Buffer and cleaved overnight at 4°C with tobacco etch virus (TEV) protease to remove the hexahistidine tags. Further purification was perfomed by size exclusion chromatography using on a 16/60 Superdex S200 column at 4°C in Gel Filtration (GF) Buffer (50 mM HEPES pH 7.5, 300 mM NaCl, 0.5 mM TCEP). Peak fractions were pooled and concentrated with 10 mM DTT for storage or immediate use. For crystallography, KEAP1_BTB-3-box_ and CUL3_NTD_ were mixed in a 1:1 molar ratio and incubated for 2 hours on ice before the size exclusion chromatography step. For biotinylation, proteins were expressed in BL21(DE3)-pRARE2 cells containing a BirA co-expression vector and media supplemented with 100 µM D-biotin. An additional 10 mL of a 10 mM stock solution of D-biotin (prepared in 10 mM Bicine, pH 8.3 and filter sterilized) was added to the cell cultures one hour before cell harvesting. Native and biotinylated protein masses were confirmed using intact mass spectrometry.

For baculoviral expression, plasmids were transformed into DH10Bac cells to generate bacmid DNA. Baculoviruses were then prepared from this using Sf9 insect cells. CUL3 and RBX1 viruses were co-infected to generate the CUL3-RBX1 complex, whereas KEAP1_BTB-BACK-Kelch_ was prepared alone. Large scale baculoviral expression was performed for 72 hours at 27°C. The harvested cells were resuspended in 40 mL binding buffer per 2L cell culture. PEI (1 mL) and protease inhibitors were added before cell lysis by sonication. After centrifugation of the lysate at 50 000 g, the supernatant was filtered and protein purified by nickel affinity and size exclusion chromatography as above.

### Crystallisation

Crystallisation was achieved at 20 °C using the sitting drop vapour diffusion method. Initially, crystals grew in 0.1 M MMT, 0.2 M ammonium chloride, 15 % PEG 3350, and diffracted to 6 Å. Further fine screening and seeding yielded plate-like crystals diffracting to 3.45 Å. The final protein complex crystallised at 9.4 mg/mL in 150 nL drops at a 1:2 ratio of protein to precipitant (20% PEG 3350, 10 % ethylene glycol, 0.2 M potassium citrate tribasic), using an adiitional 20 nL of seeds previously prepared in a similar condition. Crystals were cryoprotected in 20 % ethylene glycol in well precipitant and then vitrified in liquid nitrogen.

### Structure determination

Diffraction data were collected at the Diamond Light Source, station I03 using monochromatic radiation at wavelength 0.97626 Å. Automated diffraction data reduction was performed using xia2 3d, and the indexed, integrated, scaled and merged data was phased using Phaser-MR in Phenix [53] with a structure of KLHL11_BTB-BACK_ complexed to CUL3_NTD_ as the search model (PDB 4AP2). The molecular replacement (MR) structure solution was refined using Phenix [53] and Buster [54] with manual rebuilding with Coot [55]. Molprobity [56] was used to verify the geometrical correctness of the structure.

### Biolayer interferometry

Biolayer interferometry (BLI) performed on an Octet RED384 instrument (FortéBio) was used to determine the affinity of binding between different BTB-Kelch and CUL3 protein constructs as indicated. Biotinylated protein ligand buffered in 50 mM HEPES, 300 mM NaCl, 0.5 mM TCEP, and 10 mM DTT was used at 0.16 mg/mL to immobilise ligand onto streptavidin-coated fiber optic tips (FortéBio) to yield a binding response of 7-8 nm. Serial dilutions of the test analyte protein in the same buffer supplemented with 0.01 % TWEEN-20 were placed in the relevant wells, with matching buffer in the reference wells. Association and dissociation phases were recorded as indicated. Steady state equilibrium and kinetic fits were performed by global data analyses in the ForteBio Data Analysis 9.0 software using a 1:1 binding model.

### TR-FRET measurements

Unless otherwise noted, experiments were performed in white, 384-well microtiter plates (Corning 3572) in 30 μL assay volume. TR-FRET measurements were acquired on a Tecan SPARK plate reader with SPARKCONTROL software version V2.1 (Tecan Group Ltd.), with the following settings: 340/50 nm excitation, 490/10 nm (Tb), and 520/10 nm (AF488) emission, 100 μs delay, 400 μs integration. The 490/10 nm and 520/10 nm emission channels were acquired with a 50% mirror and a dichroic 510 mirror, respectively, using independently optimized detector gain settings unless specified otherwise. The TR-FRET ratio was taken as the 520/490 nm intensity ratio on a per-well basis.

### Protein labeling

Full-length KEAP1 (Sino Biological 11981-HNCB) and KLHL11_BTB-BACK_ were labeled with CoraFluor-1-Pfp, and CUL3_NTD_ was labeled with AF488-Tfp, as previously described [37]. The following extinction coefficients were used to calculate protein concentration and degree-of-labeling (DOL): KEAP1 *E*_280_ = 80,335 M^-1^cm^-1^, KLHL11 *E*_280_ = 34,295 M^-1^cm^-1^, CUL3_NTD_ *E*_280_ = 36,705 M^-1^cm^-1^, CoraFluor-1-Pfp *E*_340_ = 22,000 M^-1^cm^-1^, AF488-Tfp *E*_495_ = 71,000 M^-1^cm^-1^. Protein conjugates were snap-frozen in liquid nitrogen, and stored at -80ºC.

### Determination of equilibrium dissociation constant (*K*_D_) of AF488-labeled CUL3 toward CoraFluor-1-labeled KEAP1 and KLHL11

CoraFluor-1-labeled KEAP1 and KLHL11 were diluted to 1.5x (30 nM and 3 nM, respectively) in assay buffer (25 mM HEPES, 150 mM NaCl, 1 mM DTT, 0.5 mg/mL BSA, 0.005% TWEEN-20, pH 7.5). AF488-labeled CUL3_NTD_ was added in serial dilution from 3x stock solutions in assay buffer (1:2 titration, 12-point, c_max_ = 1,000 nM) and allowed to equilibrate for 2 h at room temperature before TR-FRET measurements were taken. Data were fitted to a One Site – Total Binding model in Prism 9.

### Measurement of dissociation rate constants (*k*_off_) by TR-FRET

Solutions of: (i) 20 nM CoraFluor-1-labeled KEAP1, 300 nM AF488-labeled CUL3_NTD_, and (ii) 2 nM CoraFluor-1-labeled KLHL11, 45 nM AF488-labeled CUL3_NTD_ were prepared in assay buffer and allowed to equilibrate for 2 h at room temperature before initial (*t* = 0) TR-FRET measurements were taken. Following addition of 25 μM unlabeled KLHL11, the time-dependent change of TR-FRET intensity was recorded (in 30 s intervals) over the course of 30 min. Data were normalized and fitted to a one-phase decay model in Prism 9.

### TR-FRET protein displacement assays

The following assay parameters have been used (all 1.5x): (i) 30 nM CoraFluor-1-labeled KEAP1, 300 nM AF488-labeled CUL3_NTD_ in assay buffer, (ii) 3 nM CoraFluor-1-labeled KLHL11, 45 nM AF488-labeled CUL3_NTD_ in assay buffer. In all cases, protein constructs were added in serial dilution from 3x stock solutions in assay buffer (1:3 titration, 12-point, c_max_ = 25 μM) and allowed to equilibrate for 2 h at room temperature before TR-FRET measurements were taken. The assay floor (background) was defined with the 25 μM CUL3_NTD_ dose, and the assay ceiling (top) was defined via a no-protein control. Data were background corrected, normalized, and fitted to a four-parameter dose response model [log(inhibitor) vs. response – Variable slope (four parameters)] using Prism 9, with constraints of Top = 1, and Bottom = 0.

### Calculation of protein *K*_D_ values from measured TR-FRET IC_50_

For TR-FRET protein displacement assays, we have determined the *K*_D_ of the respective fluorescently labeled protein tracer under each assay condition. Protein *K*_D_ values have been calculated using Cheng-Prusoff principles, outlined in equation 1 below:

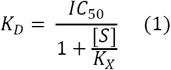

Where IC_50_ is the measured IC_50_ value, [S] is the concentration of the fluorescent protein tracer, and *K*_X_ is the *K*_D_ of the fluorescent protein tracer for a given condition [57].

## Abbreviations

BLI: biolayer interferometry
BTB: bric-à-brac tramtrack and broad complex
BACK: BTB and C-terminal Kelch
CDDO: 2-cyano-3,12-dioxooleana-1,9(11)-dien-28-oic acid
CI: confidence interval
DLG: Asp-Leu-Gly
ETGE: Glu-Thr-Gly-Glu
ITC: isothermal titration calorimetry
KEAP1: Kelch ECH associating protein 1
KLHL: Kelch-like
NRF2: Nuclear factor erythroid 2-related factor 2
Pfp: pentafluorophenyl
SCF: SKP1-CUL1-F-box
Tfp: Tetrafluorophenyl
TR-FRET: time-resolved Förster resonance energy transfer

## Data availability

The atomic coordinates and structure factors have been deposited in the Protein Data Bank, Research Collaboratory for Structural Bioinformatics, Rutgers University, New Brunswick, NJ (http://www.rcsb.org/) with PDB code 5NLB.

## Acknowledgements

The authors would like to thank Diamond Light Source for beamtime (proposal mx10619), as well as the staff of beamline I03 for assistance with crystal testing and data collection. We also thank David A Cruz Walma for help with the schematic in Fig. 1. This research was supported by the CHDI Foundation to R.J.A., S.G.B. and A.N.B. A.N.B. also acknowledges funding from the Innovative Medicines Initiative 2 Joint Undertaking (JU) under grant agreement No 875510 (EUbOPEN). The JU receives support from the European Union’s Horizon 2020 research and innovation programme and EFPIA and Ontario Institute for Cancer Research, Royal Institution for the Advancement of Learning McGill University, Kungliga Tekniska Hoegskolan, Diamond Light Source Limited. R.J.A. and A.N.B. also received support from The SGC (charity no. 1097737) which received funds from AbbVie, Bayer Pharma AG, Boehringer Ingelheim, Canada Foundation for Innovation, Eshelman Institute for Innovation, Genome Canada, Innovative Medicines Initiative (EU/EFPIA) [ULTRA-DD grant no. 115766], Janssen, Merck KGaA Darmstadt Germany, MSD, Novartis Pharma AG, Ontario Ministry of Economic Development and Innovation, Pfizer, Saõ Paulo Research Foundation-FAPESP, Takeda, and Wellcome [106169/ZZ14/Z]. This work was further supported by NSF 1830941 to R.M. N.C.P. was supported by a National Science Foundation Graduate Research Fellowship (DGE1745303).

## Author contributions

R.J.A. prepared recombinant proteins, performed the final crystallography, solved the structure and performed the biolayer interferometry. S.G.B. conducted initial crystal screens and protein production. N.C.P performed the TR-TRET experiments. R.J.A., N.C.P., R.M. and A.N.B. contributed to data analysis and manuscript preparation. R.M. and A.N.B helped with supervision and study design. All authors approved the final manuscript.

