## Supplemental Tables and Figures for "Structural and biochemical characterization establishes a detailed understanding of KEAP1-CUL3 complex assembly"

**Table S1. Diffraction data collection and refinement statistics**

| | KEAP1-CUL3 |
|---|---|
| **Data collection[a]** | |
| Beamline | Diamond I03 |
| Wavelength (Å) | 0.9763 |
| Spacegroup | C2 2 21 |
| Cell dimensions | |
| a/b/c (Å) | 41/233.38/164.71 |
| α/β/γ (°) | 90/90/90 |
| Resolution range (Å) | 40.38 - 3.45 (3.573-3.45) |
| Total reflections | 70080 (6938) |
| Unique reflections | 10902 (1049) |
| Multiplicity | 6.4 (6.6) |
| Completeness (%) | 99.55 (99.24) |
| Mean I/σI | 12.96 (1.46) |
| $R_{merge}$ | 0.08226 (1.522) |
| $R_{meas}$ | 0.08983 (1.653) |
| $R_{pim}$ | 0.03557 (0.6393) |
| CC1/2 | 0.999 (0.611) |
| **Refinement[a]** | |
| Reflections used in refinement | 10899 (1049) |
| Reflections used for $R_{free}$ | 520 (60) |
| MR model | 4AP2 |
| Copies in ASU | 1 |
| $R_{work}/R_{free}$ (%) | 23.8/28.8 (33.8/36.2) |
| CC work/CC free | 0.971/0.950 (0.788/0.526) |
| Number of non-hydrogen atoms | 3882 |
| Protein residues | 502 |
| rmsd | |
| Bond lengths (Å) | 0.011 |
| Bond angles (°) | 1.44 |
| Ramachandran favoured (%) | 92.14 |
| Ramachandran allowed (%) | 7.06 |
| Ramachandran outliers (%) | 0.81 |
| Rotamer outliers (%) | 7.87 |
| Clashscore | 8.57 |
| Average B-factor ($Å^2$) | 167.14 |
| Protein Data Bank code | 5NLB |

[a] Values in parentheses are for the highest resolution shell.

**Table S2. Equilibrium dissociation constants ($K_D$ values) determined in $KLHL11_{BTB\text{-}BACK}$ / $CUL3_{NTD}$ TR-FRET protein displacement assay.**

| Protein construct | $K_D$ (nM) | $K_D$ 95% CI (nM) |
|---|---|---|
| $KLHL11_{BTB\text{-}BACK}$ | 14 | 13-15 |
| $CUL3_{NTD}$ | 8 | 7-9 |
| $CUL3_{NTD\Delta 22}$ | 1,840 | 1,702-1,990 |
| $KEAP1_{BTB\text{-}3\text{-}box}$ | 720 | 650-797 |
| $KEAP1_{BTB\text{-}BACK\text{-}Kelch}$ | 798 | 730-872 |

**Table S3. Equilibrium dissociation constants ($K_D$ values) determined in KEAP1$_{FL}$ / CUL3$_{NTD}$ TR-FRET protein displacement assay.**

| Protein construct | $K_D$ (nM) | $K_D$ 95% CI (nM) |
|---|---|---|
| CUL3$_{NTD}$ | 129 | 117-142 |
| CUL3$_{NTD\Delta 22}$ | 13,783 | 12,047-16,119 |
| KEAP1$_{BTB\text{-}3\text{-}box}$ | 1,042 | 943-1,151 |
| KEAP1$_{BTB\text{-}BACK\text{-}Kelch}$ | 396 | 365-430 |

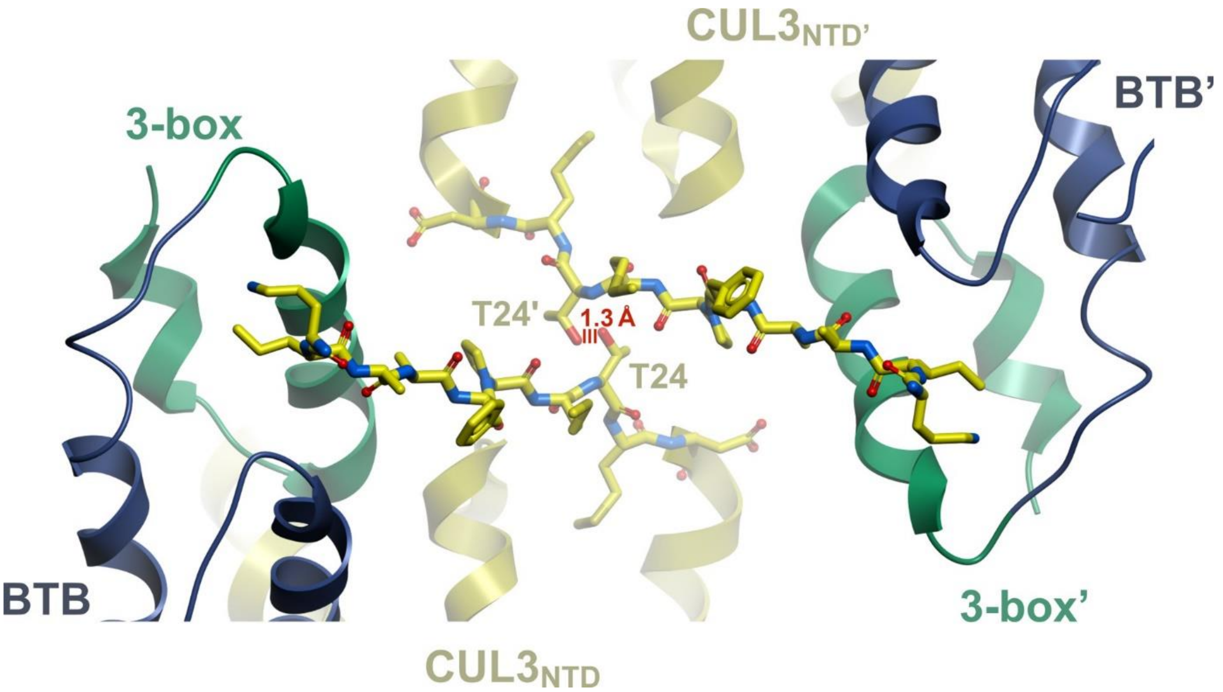


**Fig. S1. Crystal packing may create steric hindrance for the CUL3 N-terminal extension.** Two KEAP1-CUL3 complexes are shown in the crystal lattice. The $CUL3_{NTD}$ from PDB 4AP2 was modelled onto the $CUL3_{NTD}$ subunits of the KEAP1 complex to reveal the expected position of the CUL3 N-terminal extension if bound to the 3-box groove of KEAP1. This model reveals a potential clash between the N-terminal extensions in two adjacent $CUL3_{NTD}$ chains (distance 1.33 Å). Any steric hindrance could have promoted the apparent disorder in this CUL3 region.

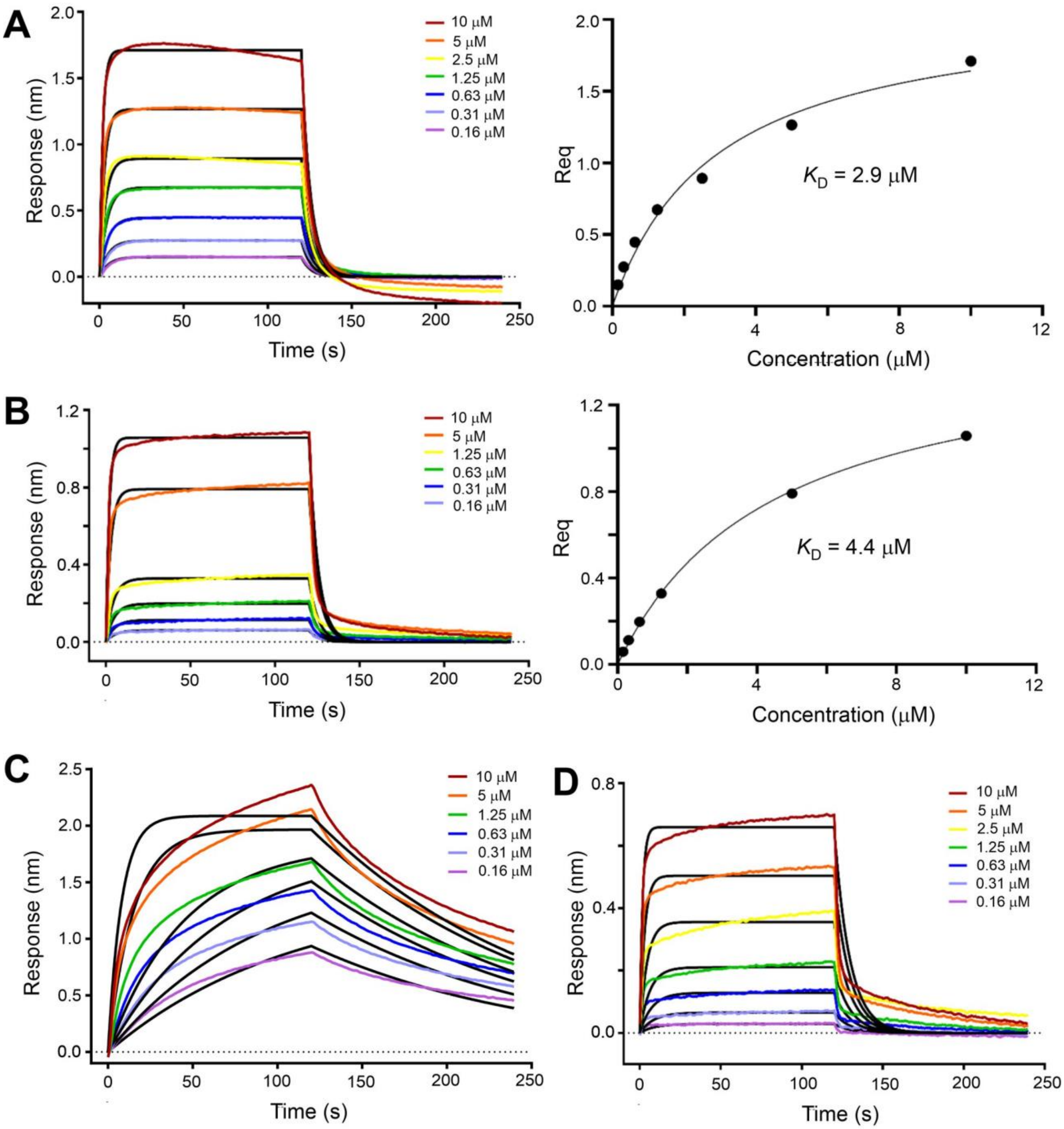


**Fig. S2. Biolayer interferometry (BLI) experiments with certain KEAP1 and CUL3 constructs showed significant deviation from a standard Langmuirian 1:1 model.** Binding measurements were recorded on an Octet RED384 instrument (FortéBio). (A) Biotinylated $KEAP1_{BTB\text{-}3\text{-}box}$ was immobilized and the binding of $CUL3_{NTD}$ showed an apparent steady state $K_D$ = 2.9 μM (95% CI 1.7-5.1 μM) and binding kinetics of $k_{on}$ = 3.66 x $10^4$ $M^{-1}s^{-1}$, $k_{off}$ = 1.89 x $10^{-1}$

$s^{-1}$, $^{App}K_D$ = 5.2 μM. (B) Biotinylated $KEAP1_{BTB\text{-}3\text{-}box}$ was immobilized and the binding of full length CUL3-RBX1 complex showed an apparent steady state $K_D$ = 4.4 μM (95% CI 3.7-5.3 μM) and binding kinetics of $k_{on}$ = 3.32 x $10^4$ $M^{-1}s^{-1}$, $k_{off}$ = 1.87 x $10^{-1}$ $s^{-1}$, $^{App}K_D$ = 5.6 μM. (C) Biotinylated $CUL3_{NTD}$ was immobilized on a streptavidin-functionalized sensor tip and binding to serial dilutions of $KEAP1_{BTB\text{-}BACK\text{-}KELCH}$ (residues 48-624) was assessed. Significant deviation from a standard Langmuirian 1:1 model was observed preventing reliable $K_D$ determination. (D) Biotinylated $KEAP1_{BTB\text{-}3\text{-}box}$ was immobilized and the binding of $CUL3_{NTD\Delta22}$ was assessed. Significant deviation from a standard Langmuirian 1:1 model was observed preventing reliable $K_D$ determination.
